# Continuous regimens of cortico-motor integration calibrate levels of arousal during emergence from anesthesia

**DOI:** 10.1101/2020.02.19.956789

**Authors:** Sijia Gao, Diany Paola Calderon

## Abstract

**Background:** Recovery to a conscious state when emerging from anesthesia requires full cortical desynchronization, initiation of movement and behavioral reactivity to sensory stimuli. However, the variety of cortical electroencephalogram (EEG) patterns associated with specific anesthetics and the paucity of behavioral descriptions during emergence from anesthesia have prevented EEG and behavior as feasible tracking methods to assess emerging from anesthesia. We propose a detailed combined analysis of motor and cortical activity to determine levels of arousal in rodents.

**Methods:** Using decreasing anesthetic concentrations, we simultaneously recorded local field potentials (LFPs) and movement in mice. We delineated cortical dynamics and sub-states during emergence from anesthesia by applying a smoothed-Z score to extract dominant frequencies from spectrogram. Then, we implemented KMeans to obtain cortical sub-states. Finally, we used density estimation and an abrupt change detection algorithm to segment cortical activity into periods. We used cortical sub-states obtained during isoflurane traces to supervise sub-states in sevoflurane and a pharmacologically induced-arousal model. This information together with examining videos were used to categorize behavior.

**Results:** We identified five cortical periods with restored motor behavior during emergence from isoflurane anesthetic. Periods of structured sub-states denoted when specific motor behaviors occurred. No significant differences were found when comparing the combined cortical features and motor behavior using isoflurane, sevoflurane and our arousal-rodent model. We describe graded regimens of cortico-motor activity during emergence from anesthesia to assess arousal levels.

**Conclusion:** We show cortical patterns denote gradual motor behaviors when emerging from anesthesia. Restoring motor behavior is a dynamic process that begins tens of minutes earlier than the righting reflex. Combining cortical activity and motor behavior unveils novel biomarkers to accurately track emerging from general anesthesia in rodents and likely other species.

## Introduction

Recovery to a conscious state in humans usually comprises full cortical desynchronization, initiation of movement and behavioral reactivity to sensory stimuli^1^. However, the combination of these parameters in rodents is rarely used when assessing arousal when emerging from anesthesia^2^ or when investigating the neuronal pathways underlying reversing anesthetic states in animal models^3-6^. A clear discrepancy arises when analyzing cortical activity^7^ and behavior among anesthetics. Some data suggests excluding these parameters. For example, the onset of movement during the emergence from anesthesia is associated with significant changes in gamma EEG frequencies (25-150 Hz)^8^. Nevertheless, increased gamma frequencies can occur during other behaviors^9,10^. Ambiguity in gamma frequency changes can occur during emergence from anesthesia^10,11^. The accepted view states that awakening in rodents correlates with individual events, such as spontaneous movement of a body part^12^, motor response after painful stimuli^8^ or righting reflex^4,6,13^ and implies that a single event can denote the restoration of motor behavior from a coma-like state. The righting reflex is considered the gold standard test to assess arousal in rodents, because vestibular inputs that sense head movement typically indicate awareness of the surroundings^2,14^. However, the righting reflex is retained in comatose-like rodents^15^, it lacks cortical involvement^16^ and persists after precollicular transections in rodents^17^. Taken together, these reports suggest the righting reflex is retained despite the absence of the telencephalon. So, we question whether this metric to distinguish arousal/recovery of consciousness from a mere brainstem reflex is accurate. To establish regaining integrative function, we propose that a combined detailed analysis of cortical activity and motor behavior is necessary.

We hypothesize that motor behavior is progressively restored through gradual changes in cortical activity to arousal/consciousness. So, we aimed to monitor cortical activity concurrently with motor activity, while mice emerged from deep anesthesia until the animals displayed organized movements. We analyzed how motor behavior evolves during emergence and defined cortical states (periods) accompanying motor behaviors. We sought to determine if these cortical states are conserved in other arousal models and if they serve as predictors of motor behavior during emergence.

## Methods

### Stereotaxic Surgery

All use of laboratory animals was consistent with the *Guide for the Care and Use of Laboratory Animals* and approved by the Weill Cornell IACUC (Protocol No. 2016-0055). C-57BL6 wildtype mice at 10 to 12 weeks old were maintained in a reverse cycle and food and water were given *ad libitum.* Mice were anesthetized with isoflurane (n=18) and sevoflurane (n=6) including males and females. Animals were anesthetized with isoflurane in an induction chamber using an initial concentration of isoflurane (3% by volume in O_2_). Eyes were protected with ophthalmic ointment. Then, animals were transferred to a stereotaxic frame, and anesthetic concentration was maintained using a nose cone. We monitored isoflurane concentration using a gas analyzer (Ohmeda 5250 RGM). The skull was fixed to the stereotaxic frame. The wound edge was infiltrated with local anesthetic (bupivacaine 0.5%) and a designed-head holder device was placed on the skull to ensure durable head restraint without the need for ear bars as in Gao et al^18^. Two small craniotomies were made to target motor cortex (AP: 1.5-2 mm ML:1.2mm DV:0.2 mm). In addition to the craniotomies, a stainless-steel reference screw was placed above the visual cortex. Craniotomies were covered with silicone for seven days and Flunixin 5 mg/Kg was administered subcutaneously.

EEG transmitter implantation (EEG unrestraint mice): animals (n=5) were induced at 3% isoflurane and maintained at a 1.25%(∼ 1MAC) vol. of isoflurane. We applied local anesthetic as described before. After opening an incision in the scalp, we created a subcutaneous pocket along the animal’s dorsal flank and placed the body of the transmitter (model F20-EET;DSI) into the pocket making sure biopotential leads were oriented cranially. Leads were placed in craniotomies made at stereotaxic coordinates targeting motor cortex. We secured leads using dental acrylic. Animals recovered for 7 days prior to experiments. We used the Ponemah V5 software from DSI to record LFP activity and temperature. Signals were downsampled to 1 kHz. All animals were used in the study.

### Monitoring

Spontaneous ventilation was maintained throughout the experiment. Respiratory rate was continuously monitored by the investigators. Temperature was maintained at approximately 37°C using a temperature regulator coupled to a rectal temperature probe (CWE Inc). We injected subcutaneous saline while the animal was deeply anesthetized to maintain adequate hydration. After securing the animal’s head within the head holder, we removed the silicone applied to the craniotomies and implanted electrodes for monitoring cortical activity. Continuous field potential in the cortex were recorded using the Plexon Omniplex System with Plexcontrol software (Plexon Inc., TX). To obtain the cortical field potential from wideband (0.2 Hz - 40 KHz), we used a causal 4th order butterworth filter to minimize phase distortion. Signals were downsampled to 1 kHz. Using standard methodology, the terminally anesthetized animal was intracardially perfused with paraformaldehyde (4%), followed by brain extraction, postfixation, microtome sectioning, and staining to confirm electrode placement.

Isoflurane anesthetic ramp: Animals were quickly induced with a concentration of isoflurane 3%. The anesthetic ramp was initiated when exposing animals to isoflurane with a starting concentration of 1MAC (1.25%). Anesthetic concentration was reduced at intervals of 0.25%. Each interval lasted for 30 min until reaching 0% anesthetic. Sevoflurane anesthetic ramp: Animals (n=6) were quickly anesthetized with a concentration of sevoflurane 5%. The anesthetic ramp was initiated when exposing mice to sevoflurane 3%. Anesthetic concentration was reduced in intervals lasting half an hour each until the gas was turn off.

### Drug Injections

Bicuculline methiodide (Tocris) was dissolved in 0.9% saline in a stock concentration of 20 mM and then diluted to yield a final concentration of 10 mM. 200 nl of drug were injected in mice (n=3) using a microinjector (Micro 4, WPI) via Hamilton syringe connected to a cannula using tygon tubing at a rate of 200 nl/15s as we previously described in Gao and colleagues^18^.

### aNGC-arousal model

We previously published a model of arousal in which we awaken animals via the anterior portion of the nucleus gigantocellularis (aNGC) from a low brain activity state(constant exposure to isoflurane 1.3%)^18^. To assess the level of arousal reached by this animal model using the proposed method in this manuscript, we implanted animals(n=3) with a unilateral cannula to acutely microinject bicuculline in aNGC. Stereotaxic coordinates: anterior-posterior (AP):-5.6 mm from bregma; (ML):0.5 mm; dorsoventral (DV)-4.25 mm. We also implanted electrodes in the motor area as described above. After animals recover for 7 days, we microinjected bicuculline in aNGC similarly and assessed motor cortical activity and behavior.

### LFP Spectral analysis

To examine changes in spectral content evolving over time, we computed spectrograms using the Thomson multitaper method implemented in the Chronux toolbox^19,20^ in Matlab (Mathworks). We used the function *mtspecgramc* to compute cortical spectrogram with the following parameters: frequency band = 2-150 Hz, tapers = [3, 5], movingwin = [5, 2.5] seconds. In Bic experiments, we used movingwin=[5, 0.5] seconds. Spectral estimates were approximately chi-squared distributed, which is skewed^21,22^. Therefore we log-transformed (converted to dB) the fractional power and then removed its median over the whole trace^23^

### Detection of dominant frequency band

We define dominant frequency band in a cortical spectrum, the band that surpasses other bands in power. It is identified by first measuring the mean power of each 50 log-spaced frequency^24^ between 2-150Hz (denoted as *x* = [*x*^1^, *x*^2^, …, *x*^50^]) and then by detecting peaks in *x* via a smoothed Z-score thresholding algorithm (stack overflow)^25^

The smoothed Z-score thresholding algorithm takes *x* as input and outputs a vector *y* = [*y*^1^, *y*^2^, …, *y*^50^], which is a sequence of “0”, “1” or “-1”. Zero represents no peak, −1 a negative peak or 1 a positive peak detected at a frequency span. In principle, peaks are identified by constructing a moving mean μ and a moving standard deviation σ from a smoothed signal *x*^*smooth*^. The algorithm requires 3 parameters to be specified: *lag* = it represents how much of the data will be smoothed. Number of last several datapoints in *x*^*smooth*^ to update μ, σ; *threshold* = deviation from μ quantified in σ to notify a peak and *influence* (ranging between 0 and 1) = influence of new datapoints on *x*^*smooth*^. In this paper, we set parameters *lag*=2, *threshold*=2 and *influence*=0.1.

The algorithm is summarized as follows: We first initialized *x*^*smooth*^ using the first *lag* number of datapoints in *x and set* μ = mean (*x*^*smooth*^), σ = std (*x*^*smooth*^).

For *j* = *lag* + 1 *to N* we did the following: If *abs*(*x*^*j*^ − μ) >*threshold* × σ, the algorithm signified a “1” (positive) or “-1” (negative). We then concatenated *x*^*smooth*^ with a new datapoint=*influence* × *x*^*j*^ +(1-*influence*) × (last element in *x*^*smooth*^). Likewise, if *abs*(*x*^*j*^ − μ) < *threshold* × σ, *x*^*smooth*^ was concatenated by *x*^*j*^. Finally, we updated μ, σ using the last *lag* number of datapoints in *x*^*smooth*^. After obtaining *y* = [*y*^1^, *y*^2^, …, *y*^50^], we first found *y*^*i*^ in 3.5-4Hz with maximum power> 4dB. We zeroed *y*^*i*^ with frequencies <4Hz to avoid artifacts often seen below this frequency and then chose all spans of “1”s with length ≥5. The span with highest mean power was determined as the single dominant frequency band.

### Classification of cortical states

We clustered span indices for dominant frequency bands in 3 animals during their emergence from isoflurane using KMeans and obtained 5 clusters: 4-8Hz (theta), 10-20Hz (alpha), 20-40Hz (beta), 30-100Hz (gamma), 70-130Hz (high gamma). The optimal number of clusters is determined using Elbow method on inertia (Fig. S1; wiki on elbow, inertia from python sklearn)^26^. To classify cortical states, we assigned span index of dominant frequency bands to its nearest centroid. However, we used the power in 3.5-4 Hz to exclusively characterize period 2 as this feature is remarkably specific and consistent across all conditions in which period 2 appeared.

### Segmentation of cortical period

We delineated cortical dynamics through occurrence density of the classified cortical states and then segmented it into periods by applying an abrupt change detection algorithm. To achieve density estimation, we implemented the locfit.m function in the Chronux toolbox in MATLAB. In principle, time instants of a cortical state *i* behave as single action potentials of a neuron. In the locfit.m function, we chose density estimation-type family=“rate” and “nearest neighbor” smoothing method with parameter 0.05. We interpolated locfit output using interp1.m function and applied z-score normalization to get the normalized density per cortical state per second. We segmented by first obtaining a matrix *A* through concatenating normalized density of all cortical states (3.5-4Hz, 4-8Hz, 10-20Hz, 20-40Hz, 30-100Hz and 70-130Hz). Cortical states with number of time instants <100 are discarded. We then implemented the findchangepts.m function (from Matlab) to find indices where local mean of *A* changed most dramatically through minimizing sum of residual error of each segmented region. We specified parameters “MaxNumChanges” = 8, “MinDistance” = 600 for long ramp exp and “MinDistance” = 60 for short ramp, Bic exp. Cortical periods were finally segmented based on these detected changepoints in a manual fashion.

### Computation of transition matrix for cortical periods

To determine whether brain activity is sequentially ordered in periods when emerging from anesthesia, we computed a transition matrix for cortical periods in isoflurane (n=9 animals), sevoflurane (n=6 animals) and isoflurane short ramp (n=3 animals). Element located at *i*th row, *j*th column in the transition matrix represents total number of transitions from period *i* to *j* observed on samples. We normalized rows in the transition matrix so that each row added up to one.

### Motor behavior observed from the video

Motor behavior was visually inspected by an investigator blind to the experimental procedure. Motor behavior has been classified as follows:

1. Trunk movement.
2. Hindlimb movement. It includes two type of movements:
  a. abduction, adduction while animals lay down
  b. limb alternation
3. Body Wobbling-Weak weight bearing:
  a. weak weight bearing with limb retraction
  b. weak weight bearing with dragging of limbs
  c. Body quivering-Abrupt and short whole-body movements
4. Organized Movement:
  a. Full limb retraction with wide stand
  b. full weight bearing (touchdown)
  c. Jumping

### Movement detection using a vibration sensor

Animal movements were detected by using a two centimeter-piezo element sensitive to vibration placed below the animal’s body while animals emerged from anesthesia. LFPs signals and voltage changes as a result of vibration were simultaneously recorded using the Plexon OmniPlex system. A video camera was synchronized to LFP recordings to observe an animal’s behavior. We removed offset in sensor signal every 1s without overlap.

### Sample sizes

To determine the sample sizes of the experimental groups we performed pilot experiments with 3 mice for the pharmacologic experiments^18^ and for those mice exposed to prolonged anesthetic ramps. We considered the strength of the effect and the variance across the groups to determine the sample size (number of units). For experiments in which we assessed motor arousal, we estimated sample sizes using data previously published^12,27,28^. We estimated ideal samples by conducting power analysis. All experiments met or exceeded the ideal sample size.

### Statistical analyses for experiments

To perform anesthetic ramps C-57/BL6 mice were randomly assigned to isoflurane, sevoflurane and aNGC-arousal model groups. All experimental animals were subjected to the same surgical procedures.

### Code availability

The function custom-written in Matlab to detect dominant frequency ranges during emergence from isoflurane in mice is available from the corresponding author.

## Results

### Cortical features coincide with restoring motor behavior during the emergence from anesthesia

To determine whether restoring movement in rodents during emergence is associated with reliable signals detected by LFP, we anesthetized C-57/BL6 mice (n=18; males & females) using isoflurane at a concentration of 3% vol. and maintained animals at a concentration of 1.25% vol. for 30 minutes to reach an initial stable concentration. During this time, animals were head restrained using a head holder previously implanted (see methods) (**Fig 1c**). We monitored the motor cortex using bilateral LFPs during the reduction of anesthetic (**Fig 1a)** and up to 30 minutes after turning off the volatile gas. This time is sufficient to fully expel the anesthetic from the animal’s body^29^. We detected animal movements by using a two centimeter-piezo element sensitive to vibration placed below the animal’s body while animals emerged from anesthesia to determine the strength and occurrence density of the movements (mild: low voltage change; strong: high voltage change). We simultaneously recorded LFPs signals and voltage changes as a result of vibration using the Plexon OmniPlex system. We also synchronized a video camera with LFP recordings to visualize and examine animal’s behavior. Given the variability in the power of the different frequencies evidenced in the cortical spectrogram while reducing anesthetic in a stepwise fashion (**Fig 1b;** top panel), we applied a smoothing Z-score thresholding algorithm to extract frequencies (**Fig 1b**; middle panel) that occupied most power (two-dimension vector comprises lower and upper limit frequencies; see methods). Then, we determined the presence of possible cortical sub-states through clustering this two-dimension vector. We obtained five clusters of major frequency ranges (**Fig 1b;** bottom panel). A first cluster (gray) corresponded to frequencies ranging between 4-8 Hz. A second cluster (black) included frequencies between 10-20 Hz, while a third cluster (blue) contained frequencies (20-40 Hz). The purple cluster included higher frequencies corresponding to 30-100 Hz and the lilac corresponded to the frequency range of 70-130 Hz. We examined the goodness of the resulting clusters using the sum of squared distances of samples to their closest centroid (inertia) (**Supplementary Fig 1**).

**Fig. 1.**
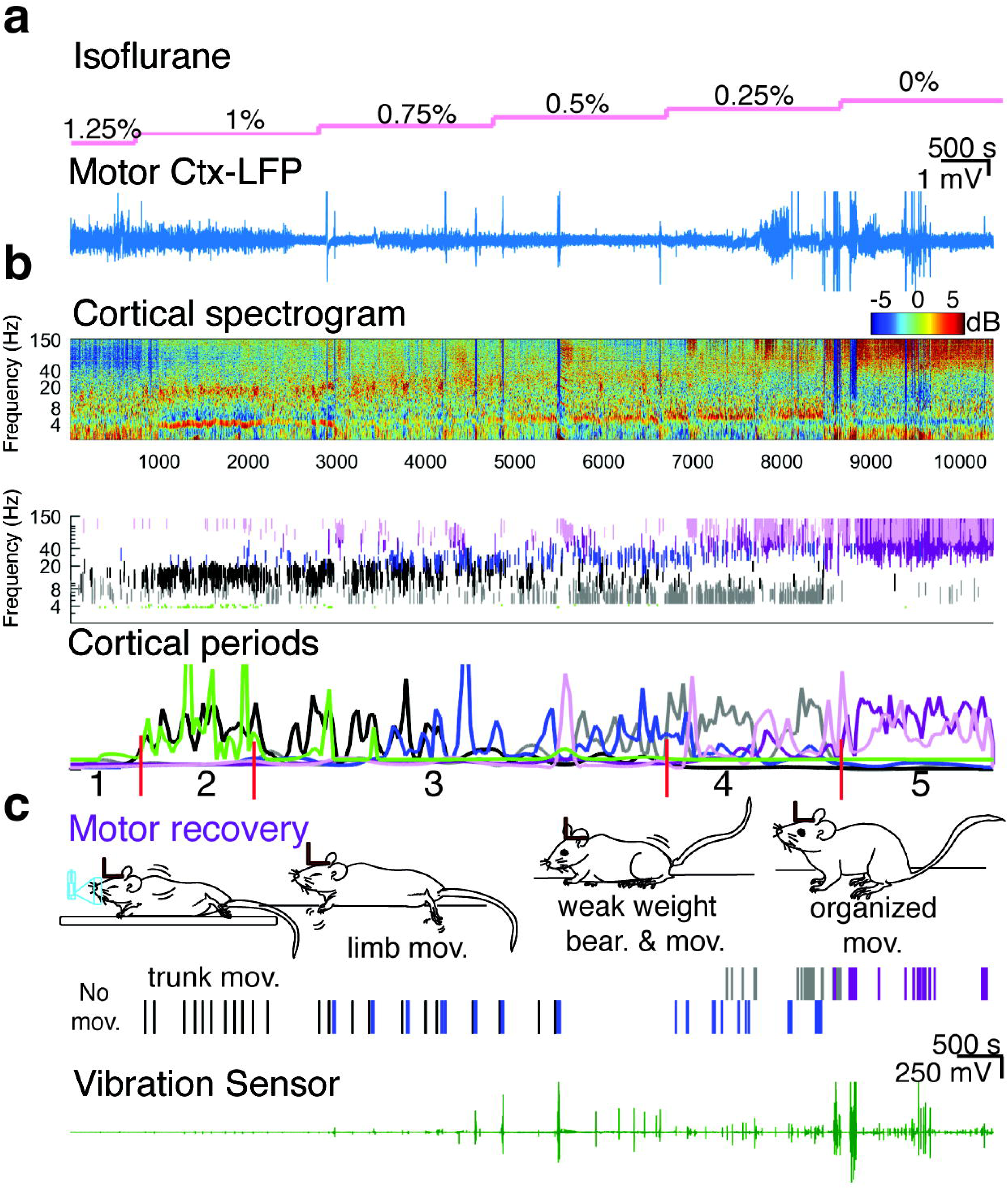
Cortical features define restoring motor behavior during emergence from anesthesia. (**a**) Representative traces of raw LFP recorded in motor cortex during exposure to isoflurane in mice (pink line). We ramped down the concentration of anesthetic from 1.25% (1MAC) to 0% at intervals of 0.25% each. (**b**) Top: Normalized spectrogram (deviation from median). Color bar shows power in decibels. Medium panel shows the dominant frequency intervals: green dots (3.5-4 Hz), gray(4-8 Hz), black(10-20 Hz), blue(20-40 Hz, purple(30-100 Hz) and lilac(70-130 Hz) after clustering prominent frequencies. Bottom: Cortical segmentation by applying the density estimation function and an abrupt change detection algorithm (periods 1-5). (**c**) Schematic depicts the evolution of movements observed in the video and detected by the sensor (green trace) during the emergence from anesthesia. Animals showed trunk movements (black lines), forelimb and hindlimb movements (blue lines), weak weight bearing (WWB) posture (gray lines), together with limb movements and organized movements (purple lines).

When animals were exposed to isoflurane 1.25% (1 MAC), (**Fig 1b** (representative trace) we observed the dominance of theta (4-8 Hz) and intermittent occurrence of high gamma (70-130Hz). During this period, mice remained immobile. We defined this condition as period 1. However, when reducing the concentration of isoflurane, we observed a state with a noticeably increased power at the 3.5-4 Hz (green) and 10-20 Hz band (black). During this cortical activity pattern, we detected brisk trunk movements as the prevalent behavior (distinguishable from breaths). We defined this condition as period 2. Although the 3.5-4 Hz was not obtained through the clustering, we added it as part of the characterization of this period given its consistency across animals and exclusive presence during the second period. As we ramped down the anesthetic (0.75%-0.5%), we saw a characteristic predominance of the 20-40 Hz band (blue) over the rest of the frequencies accompanied by chirps of 30-100 Hz (purple) and 70-130Hz (lilac). This cortical activity aligned with the natural progression of movements. In addition to trunk movements, we observed that forelimb and hindlimb movements progressed from mild to strong, including limb abduction and adduction (**Fig 1c & 2a**) and limb alternations. On occasion, the disorganized limb movements resulted in a posture where the animals temporarily and weakly held its weight on limbs. We defined the combination of this cortical and motor activity as period 3. Then, animals transitioned to a state in which the 30-100 Hz and the 70-130 Hz band increasingly dominated together with the lively presence of the 4-8 Hz band (Period 4; **Fig1b and 2a**). During this cortical pattern, the posture evolved to a persistent partial support of their body weight (squatting position; **Fig 1c**). Combination of simultaneous movement of multiple body parts derived in generalized body movements. Interestingly, when the 4-8 Hz power dominated in this period, the animals significantly reduced movement. Finally, when gamma frequencies predominated (30-100Hz & 70-130 Hz) and the alpha and beta bands significantly decreased, animals showed organized movements that evolved from a wide stand to full weight-bearing posture and active jumping. This period persisted until we turned off the anesthetic and finished the recording. Note that detecting cortical features was not linked to the nominal value of the anesthetic concentration (**Fig 1a-b**).

To examine whether the sequence of movements observed in head restrained mice was similar to those found in unrestrained conditions, we examined cortical activity and motor behavior in animals previously implanted with a wireless EEG transmitter (Data Science International). The transmitters were previously implanted in the back of the animals. Two leads of the transmitter were extended to the skull to bilaterally record motor cortical activity. Animals were placed in a gas tight chamber, and isoflurane was delivered using a calibrated vaporizer in 50% air/50% O2. Anesthesia was induced with 3% isoflurane at a flow rate of 8L/min for 2 min. The anesthetic ramp (1.25-0%) was provided using a flow rate of 2L/min. In addition to the cortical and behavioral changes reported in head restraint mice (**Fig 1&2a**), we noticed distinguishable head movements minutes after the initial trunk movements originated (**Fig 3c)**. Cortical changes and progress in motor recovery were surprisingly similar to restrained mice in periods 1-5. We found no significant differences for cortical period (*F(3,111)* = 2.62, *p=*0.054) and movements(*F(3,111)* = 0.04, *p=*0.98; Three-way ANOVA) (**Fig 2**). In some cases, we restricted the time for recording in period 5 as animals were fully active and risked chewing the temperature regulator cable.

**Fig. 2.**
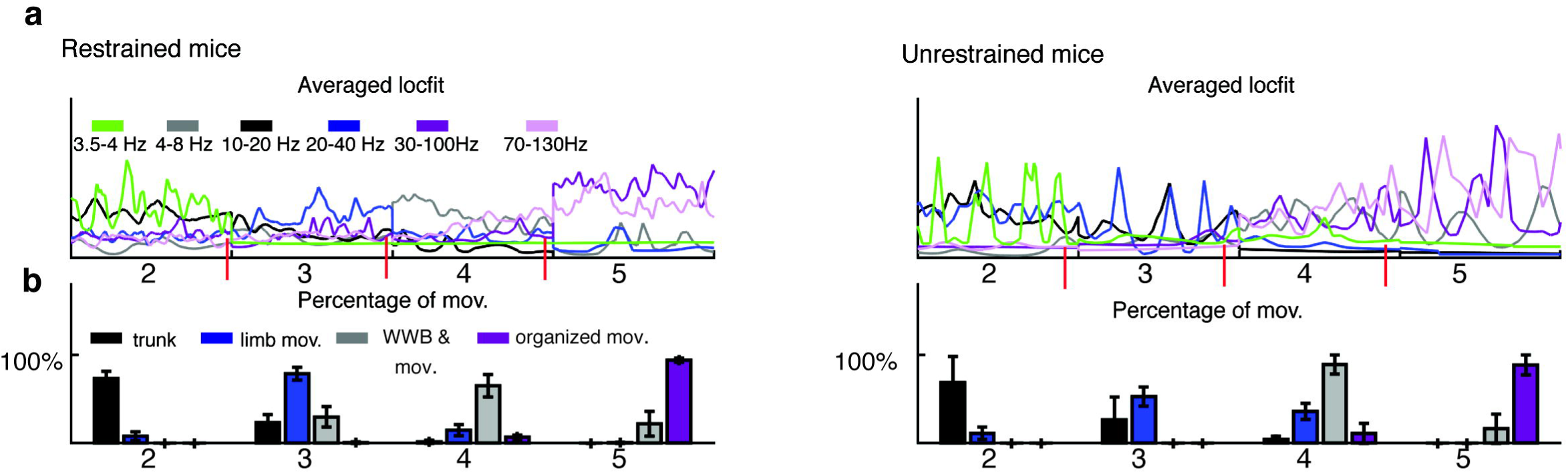
Distribution of motor behavior despite initial posture during emergence from anesthesia. (**a**) Average dominant frequency intervals and description of cortical sub-states after clustering data using kMeans and applying a density estimation to segment cortical activity into periods across multiple animals with the head restraint (n=9) or unrestraint (n=3). (**b**) Average percentage distribution of detected motor behavior per cortical periods during emergence from anesthesia in restrained and unrestrained mice. Note: We show the distribution of movement from period 2-5, as animals remained immobile in period 1.

**Fig.3.**
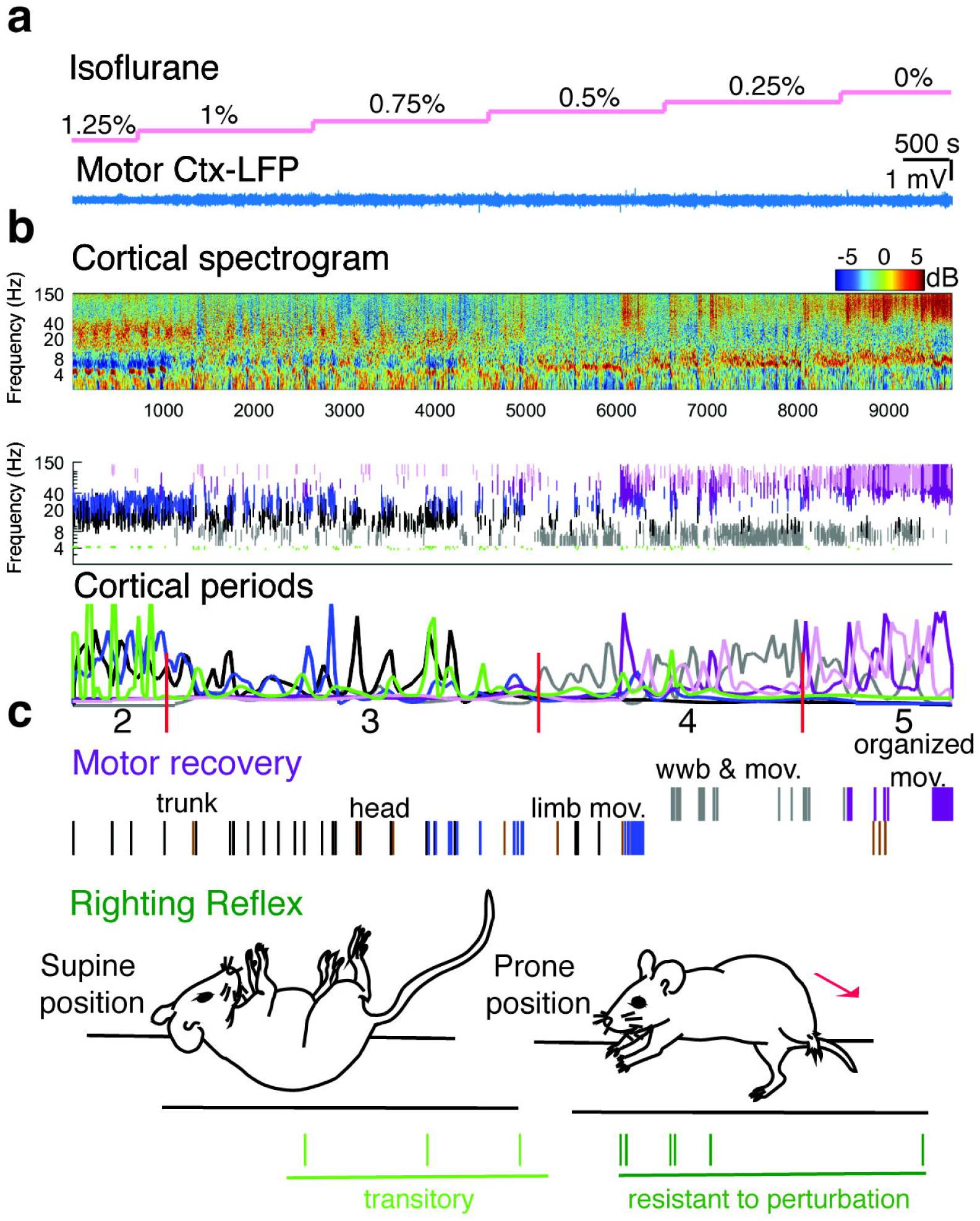
Return of the righting reflex during cortical periods and motor recovery observed at emergence. (**a**) Raw trace of LFP recorded in motor cortex during exposure to an anesthetic ramp of isoflurane (pink line). (**b**) Top: Normalized spectrogram with dominant frequency intervals obtained after clustering prominent frequencies (kMeans/smoothed-Z score algorithm). Bottom: Cortical segmentation obtained by applying the density estimation function and an abrupt change detection algorithm (periods 2-5). (**c**) Recovery of motor behavior and the appearance of two types of righting reflexes. A transitory righting reflex associated with low cortical activity, and a righting reflex resistant to perturbation occurring at higher cortical arousal.

Overall, the described periods combining cortical and motor behavior were consistently observed across all animals exposed to isoflurane ramps (restrained and unrestrained). Based on these results, we conclude that animals emerging from isoflurane anesthesia share a common dynamic process of cortical and motor arousal. Importantly, motor and cortical features detected in these long-lasting isoflurane ramps also presented when shortening the ramp by instantaneously switching concentrations from 1.25-0% isoflurane vol. (**Supplementary Fig 2**). These results indicate that these cortical and behavioral events are trackable at different rates after discontinuing the administration of anesthetic.

### Restoring motor behavior is a dynamic process beginning tens of minutes earlier than the righting reflex

Since the righting reflex is the conventional metric for arousal, we sought to establish the relationship between the return of the righting reflex and the cortical periods and motor recovery observed during isoflurane anesthesia. We used animals previously implanted with a transmitter and obtained EEG activity using the wireless telemetry system from DSI. For these experiments, we placed animals in a supine position as described by others^17,30^ and shown in the schematic (**Fig 3c**). Animals were subjected to the same anesthetic ramp used in head-restraint mice.

Tens of minutes after the initial trunk movements (48.63+/- 10.4 standard err.), animals began to recover the righting reflex. A combination of rocking the trunk to the right and left side together with stretching the limbs resulted in rotation of the body, in which the four limbs touched the ground (**Fig 3c**). Understanding that the righting reflex is a measure to assess arousal, we expected that animals would persistently hold the righting reflex despite external perturbation. However, when we shook the chamber during periods 2 and 3 (1-0.5% isoflurane vol.), animals easily returned to a supine position (period 2: 16% of the perturbations; period 3: 75%, and period 4:8%). These results suggest that during the time when theta, alpha and beta frequencies predominated, the righting reflex was transitory (**Fig 3**). Through this period, animals showed multiple events of the righting reflex, which were easily reversed (**Fig 3c**). Conversely, when we ramped down anesthetic and reached period 4 &5 (0.25-0%), animals manifested a righting reflex resistant to perturbation (**Fig3c**), despite shaking the chamber. These results showed that the righting reflex occurred tens of minutes after the onset of movement. Moreover, the righting reflex occurred spontaneously at different anesthetic concentrations. This reflex was reversible when accompanied by features of a low-cortical activity state. In contrast, once gamma frequencies begin to predominate (period 4), the reflex persisted despite perturbation.

### Emergence from sevoflurane shares the equivalent cortical activity and motor behavior as isoflurane

Although we found a common dynamic process of motor arousal and cortical changes in all animals exposed to isoflurane, we questioned whether these cortical and behavioral features also occurred using an inhaled anesthetic with different biophysical modulation than isoflurane^31^. So, we exposed animals to sevoflurane and examined cortical activity and the corresponding motor behavior. Similar to extracting cortical features described in isoflurane, the resultant cortical periods closely resembled those seen in animals emerging from isoflurane (**Fig 4c&2a).** We found no significant differences for cortical period ((*F(3,154)* = 2.33, *p*=0.07) and movements ((*F(3,154)* = 0.15, *p*=0.92); Three-way ANOVA) between sevoflurane and isoflurane. Likewise, the progression of motor recovery showed a similar distribution among the established cortical features (**Fig 4d).** These findings suggest that the sequence of cortical regimens is indistinguishable between sevoflurane and isoflurane (**Fig 2a&4d**), even though cortical physiology responses may differ slightly between anesthetics with similar MAC concentration (**Fig 1b&4c**).

**Fig.4.**
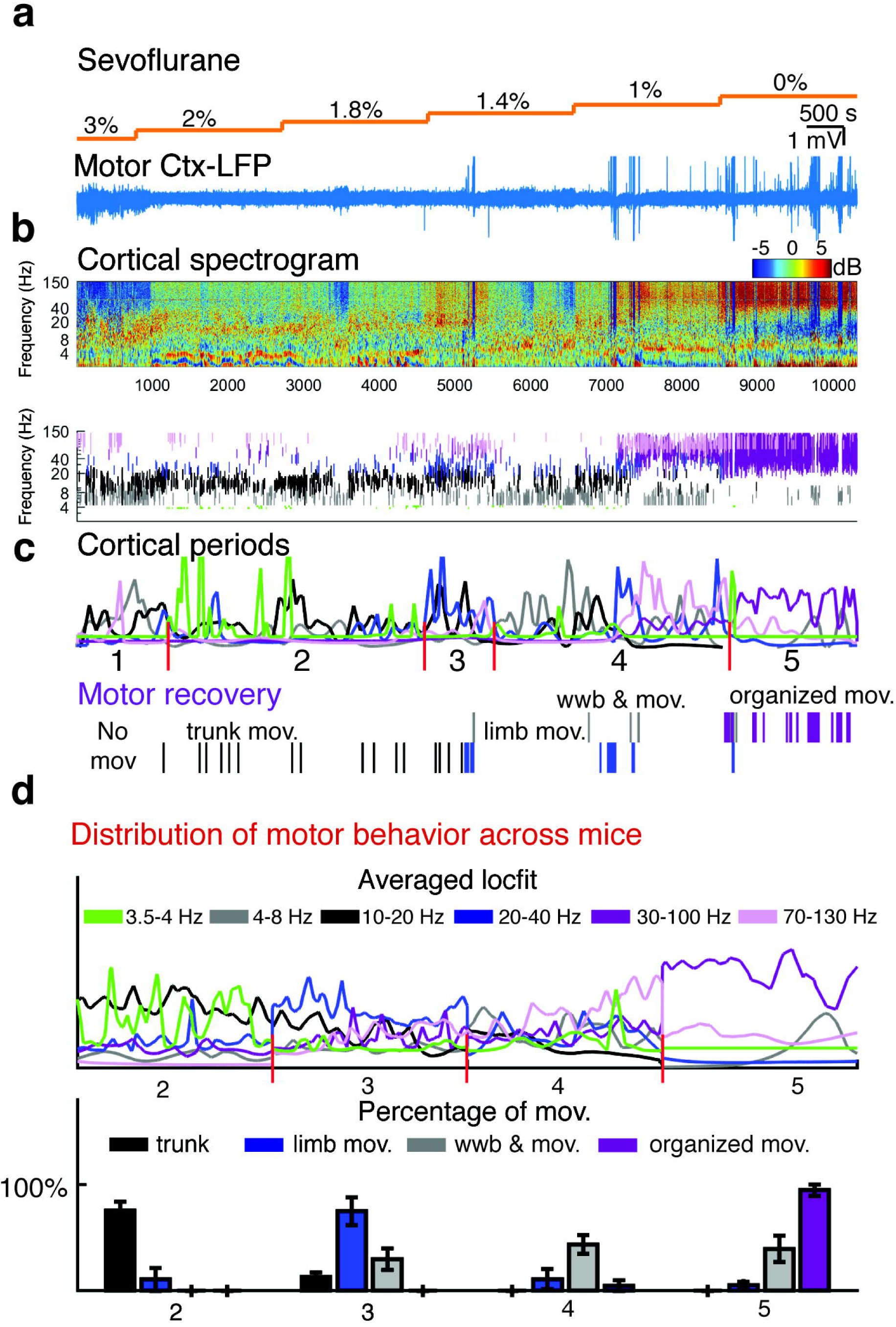
Common cortical and motor arousal features in animals emerging from sevoflurane and isoflurane anesthesia. (**a**) LFP recorded in motor cortex while exposing a mouse to an anesthetic ramp of sevoflurane (orange line). (**b**) Top: Cortical spectrograms and segmented periods after applying kMeans using the centroids obtained with isoflurane and density estimation function for a single animal. Bottom: Motor recovery from sevoflurane. (**d**) Average dominant frequency intervals and description of cortical sub-states after clustering data using kMeans and applying a density estimation to segment cortical activity into periods across multiple animals (n=6) and the average percentage distribution of the detected motor behavior per cortical periods.

### Combined motor-cortical analysis tracks arousal in a pharmacologic induced arousal model

While modulating neuronal activity in the anterior portion of the nucleus gigantocellularis (aNGC) of rodents continuously exposed to a surgical level of anesthetic, we observed robust cortical activity together with motor behavior evolving from subtle movements to more organized movements that produced grooming behavior and responsivity to external stimuli^18^. Using our proposed combined analysis, we sought to determine the level of arousal reached when stimulating aNGC in our animals. So, we exposed mice to a stable concentration of isoflurane (1 MAC) and then microinjected bicuculline (10 mM) to induce arousal via aNGC^18^. Seconds after bicuculline injection, we noticed a quick switch from period 1 to period 3 accompanied by initial brisk trunk movements that transitioned to forelimb and hindlimb movements including limb abduction and adduction. These movements aligned with those characteristic of period 3. Once there was a prominent presence of the 30-100 and 70-130 Hz sub-states, we began to observe mice bearing their weight and showed active limb movements as reported before^18^. Despite the fast arousal (10 times faster than anesthetic ramp), we observed similar cortical and behavioral characteristics. Moreover, distributing motor behavior with respect to cortical changes was comparable to results obtained by using a short anesthetic ramp (**Supplementary Fig2**). Animals temporarily restored the righting prior to entering a gamma episode as found under isoflurane. Our results identified well-defined cortical periods (**Fig 5c**) that preserved both rapid arousal and the sequence of cortical activity periods.

**Fig.5.**
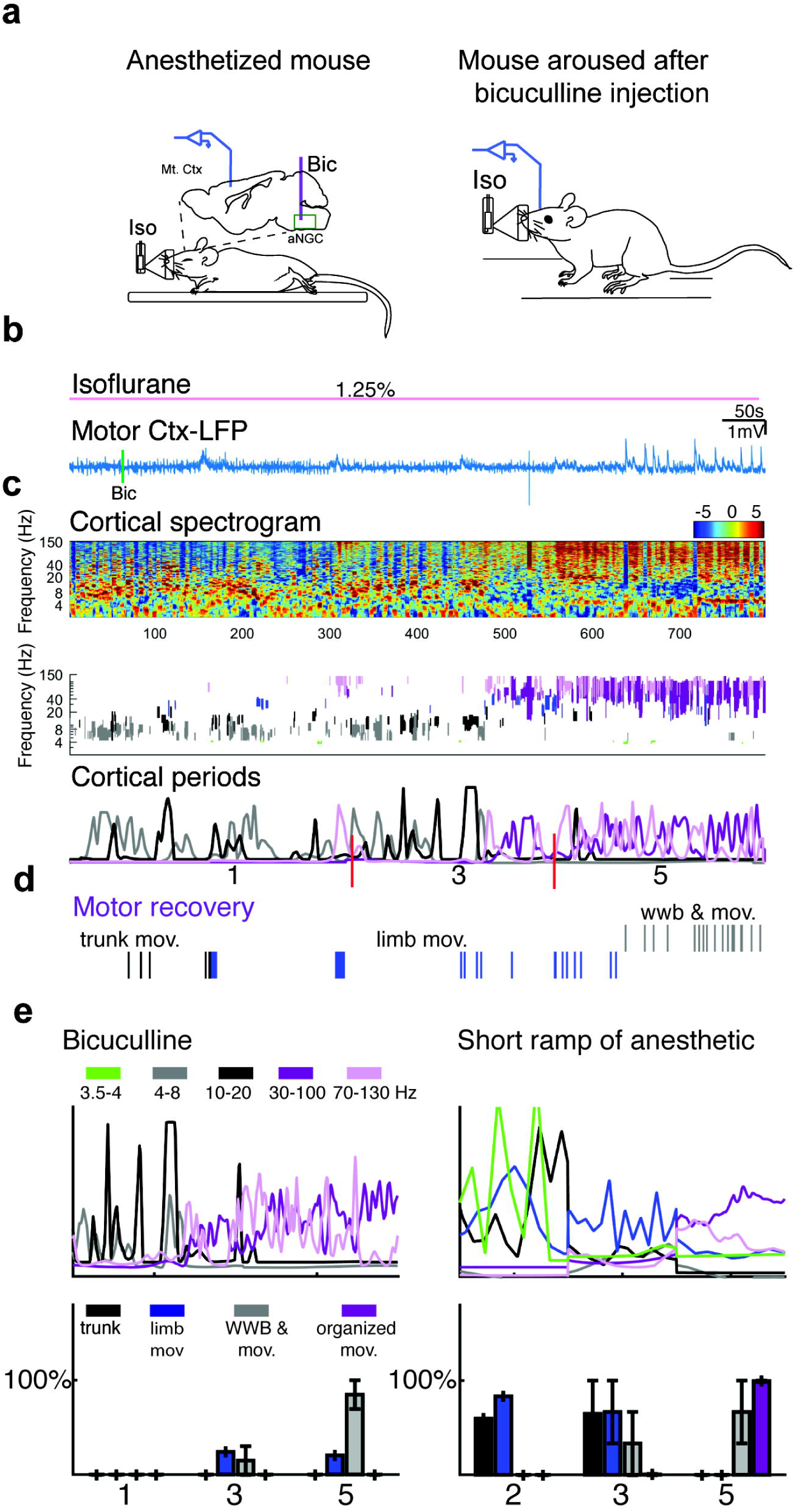
Calibration of awakening in a pharmacologic-induced arousal model. (**a**) Schematic depicts an animal deeply anesthetized and aroused after bicuculline (Bic) microinjection. Inset: Electrode location in Motor cortex (Mt. Ctx). (**b**) Cortical LFP and normalized power spectrogram before and after Bic (green line). (**c**) Dominant frequency intervals and description of cortical sub-states in a single animal. (**d**) Restoration of motor behavior during arousal after aNGC-stimulation. (**e**) Average percentage of distribution of detected motor behavior per cortical period in our aNGC-arousal model(n=3) and a short ramp of isoflurane anesthetic(n=3).

In general, the cortical periods identified here are linked to the regression across both lower and higher brain functions. In addition, they are consistently sequenced from deep to light anesthesia (**Fig 6a-c**) regardless of the prior exposure conditions.

**Fig.6.**
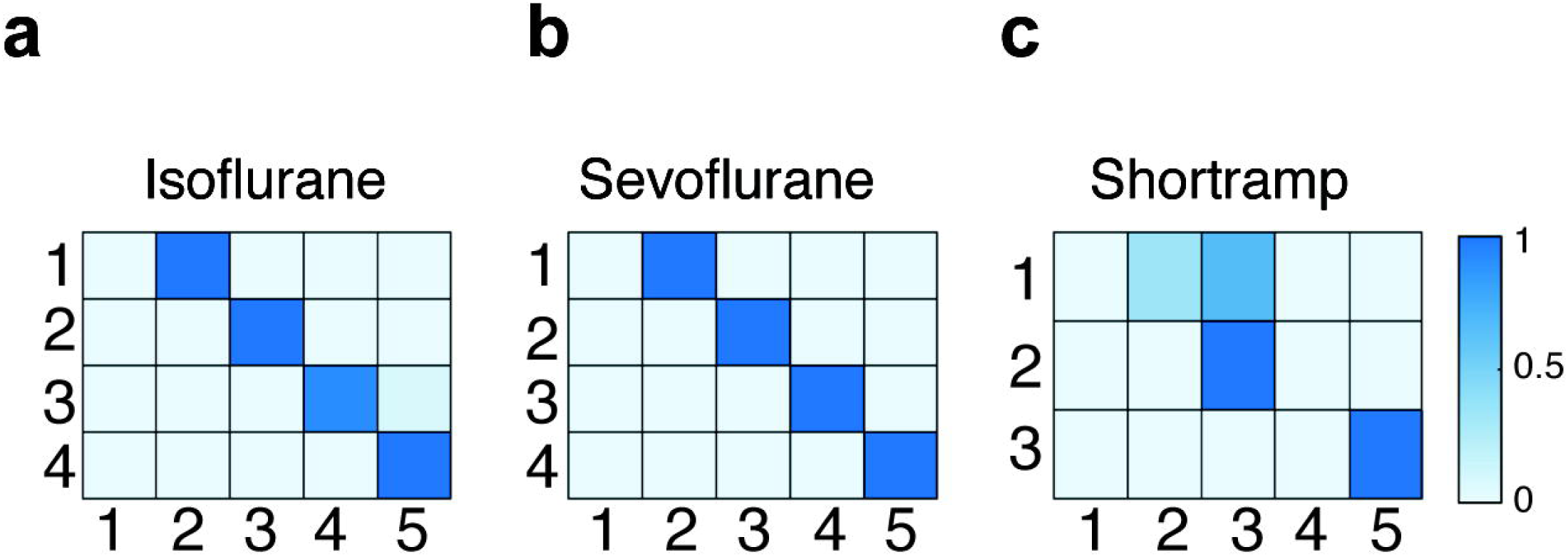
Sequence of cortical periods while emerging from anesthesia. Transition matrix for cortical periods during the emergence from anesthesia during prolonged anesthetic ramps of (**a**) isoflurane (n=9), (**b**) sevoflurane (n=6), and (**c**) fast rate of discontinuing administration of isoflurane anesthetic (n=3). The transition matrix was normalized to reach one as a total.

## Discussion

Here, we identify periods of cortical activity with common features that coincide with restoring motor behavior during the emergence from different halogenated inhaled anesthetics and a rodent model of arousal. Combining cortical activity and behavioral analyses reveals that restoring motor behavior is a dynamic process, not a single event, that begins tens of minutes before the righting reflex. Our study describes cortical periods that comprise patterned dominant-frequency bands that precisely capture shifts in the functional recovery of movement during emerging from anesthesia or arousal. Our approach reveals a methodology to determine when integrative function has been regained.

Different published methods for monitoring anesthesia in rodents mainly utilize reflexes, such as response to painful stimuli, the righting reflex^5,8,13^ and behavioral scales^32,33^. However, they are limited by the inability to assess subjects integrating the qualities that constitute the arousal process^34^. The group of Esteves and colleagues^32^ created a scale based on objective behavioral changes describing the sequences of motor behaviors during anesthesia induction and emergence. Despite using injected anesthetics (ketamine/dexmedetomidine) to anesthetize mice and atipamezole for emergence, they observed similar behaviors as the ones described here, suggesting that arousal behavior is common among inhaled and injected anesthetics. Unlike our study, their EEG analysis was limited to 7 s before and after scoring behavior, which prevented continuity of the EEG data. Since they grouped average power in wider frequencies bands, they observed an overall gradual decrease in the power of 4-12 Hz during emergence. In contrast, we observed a distinct pattern when the 4-8 Hz band increased in power at period 4 (equivalent to score 1-2 based on the described behavior) and the 10-20 Hz band decreased in power at period 4 & 5 (equivalent to score 0). We speculate that these data highlight the dynamics of different bands during particular cortical periods.

A study from Mansouri and others^33^ compared a sequence of behaviors in rats including sigh, forelimb movement, eye blinking, mastication, neck movements and the righting reflex. Although they did not find significant differences in the pattern of emergence between male and females nor the latency to the first behaviors, they found slight differences in the order of behavior presentation between propofol and isoflurane. In our study, we did not observe significant differences among gender or latency, even though our behavioral markers were more complex than those described in Mansouri’s work. Certainly, we pooled female and male data to analyze the information. The fact that the described cortical changes and progressive movements were surprisingly preserved among animals with different arousal conditions suggest that the evolution of cortical activity and movements is deterministic not random.

Interestingly, spontaneous body movements in the mouse embryo are remarkably similar to those seen during the emergence from anesthesia and the aNGC-induced arousal model. During the prenatal period, fetal movements transition from simple twitching of the trunk and limbs to complex lateral abduction and extension of limbs reaching a level of coordinated movements (limbs alternation)^35^. These movements represent the caudal to rostral maturation seen in rodents and humans that allow the fetus to respond to vibroacoustic stimuli delivered to the maternal abdomen^36^.

Moreover, we suspect that the nervous system reverses a state of anesthesia and a coma state in humans^1^ following the same organization. Thus, identifying cortical and motor features, as well as their interaction, will permit a greater understand how the brain recovers from different anesthetic-like perturbations.

Recovery of movement and cortical activity reflects restoring motor circuits, including primary motor tracts, basal ganglia, reticulo spinal tract and spinal cord^1^. Our observations indicate that the recovery is independent of the rate of arousal. We observed indistinguishable cortical periods and motor behavior in a model when mice arouse 10 times faster (short ramps and our aNGC-arousal model) than the prolonged ramps of anesthesia. Hence, cortical periods and motor behavior are identifiable, despite the rate of awakening. Although we subjected mice to different postures (prone and supine), the behavioral activity patterns remained. We found minor differences in the number of spontaneous limb movements in a supine rather than the prone posture, similar to Mendez and colleagues^37^.

By definition, anesthesia induction in mice occurs at the drug concentration when the righting reflex is lost, whereas emergence from anesthesia occurs at the concentration when the righting reflex returns^2,38^. For instance, the EC50 for the induction of halothane anesthesia is more than 2.5 times the EC50 for emergence. However, we question using the righting reflex as the appropriate metric to determine the EC50 in rodents. Our results indicate that the righting reflex may not be a proper measurement to assess arousal. During emergence from anesthesia, we noticed spontaneous, but transitory righting reflex episodes, accompanied by low cortical activity (**Fig 3**). In contrast, the righting reflex at lower anesthetic concentrations resisted perturbations and occurred with prominent gamma oscillations. We note that there was no particular signature of cortical activity associated with the righting reflex. Since the righting reflex varies during the emergence from anesthesia, it is difficult to assess the onset of awakening using this metric. For instance, several groups^3-6,33,39^ have used the righting reflex as a surrogate for arousal. However, righting reflex alone seems unspecific.

The righting reflex at Pal’s study^13^ occurs at a cortical state that shows an increase of the theta/delta ratio compared to control. Based on our assessment, this cortical feature can be found in period 3. This is a period in which the righting reflex mostly remains transitory. We obtained similar results in our aNGC-arousal model questioning the level of arousal reached by our model as well. When using our proposed combined analysis, we could accurately calibrate levels of arousal from our model and others. Studies seeking to reverse anesthesia should assess cortical activity and behavior beyond the righting reflex to identify authentic arousal. Future studies will address this question and whether this combined analysis is applicable to other type of anesthetics. Although we could outline cortical periods using a semi-automated segmentation approach (density estimation and an abrupt change detection algorithm), there were cases in which defining the critical periods was particularly challenging. Thus, our results emphasize using additional measures, such as the vibration sensor and behavior, were crucial to improve segmentation.

## Conclusions

Our study demonstrates that combining cortical activity and behavior can accurately track emergence from anesthesia. Understanding the cortical features associated with movement dynamics during general anesthesia reveals biomarkers that can inform and monitor recovery from anesthesia or muscle relaxants. The findings presented here promote using effective anesthesia and avoiding side effects from using high/low anesthetic concentrations. Moreover, these cortical and motor patterns may serve as a metric to not only reverse pharmacologically-induced coma and general anesthesia, but also examine recovery in disorders of consciousness in multiple animal models.

## Acknowledgments

We thank Hilario H., Valle C. and Islam Md. S. for helping to monitoring anesthesia ramps.

## Author Contributions

D.P.C. designed, performed experiments and analyzed experiments. SG performed data analysis of all figures and participated in some of the experiments. D.P.C., S.G. wrote the paper.

## Data availability

The data that support the findings of this study are available from the corresponding author upon reasonable request.

Correspondence and requests for materials should be addressed to

