## Supplementary Data for "Continuous regimens of cortico-motor integration calibrate levels of arousal during emergence from anesthesia"

### Supplemental information

Figure legends
Extended data Figures 1-2

**Supplementary Figures**

**Supplementary Figure 1 Determination of optimal number of clusters for kMeans clustering analysis.** To determine the optimal number of clusters, we used the Elbow method to establish the number of clusters obtained from cortical activity of animals emerging from (**a**) isoflurane and (**b**) sevoflurane. Inertia was computed as the sum of squared distances between samples and their closest centroids.

**Supplementary Figure 2 Cortical periods and motor recovery prevail during a fast rate of discontinuing isoflurane. (a)** LFP trace recorded in motor cortex while exposing a mouse to a short anesthetic ramp of isoflurane (1.25 -0 % vol.). (**b**) Top: Normalized spectrogram and dominant frequency intervals obtained after clustering dominant frequencies. Bottom: Cortical segmentation obtained by applying the density estimation function and an abrupt change detection algorithm (periods 1-5). (**c**) Recovery of motor behavior after a rapid emergence from anesthesia. Vibration sensor detected limb movements during animal laydown and during weak weight bearing posture.

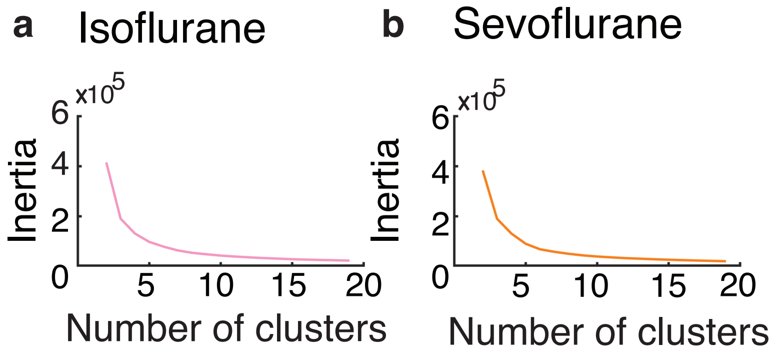


**Supplementary Figure 1**

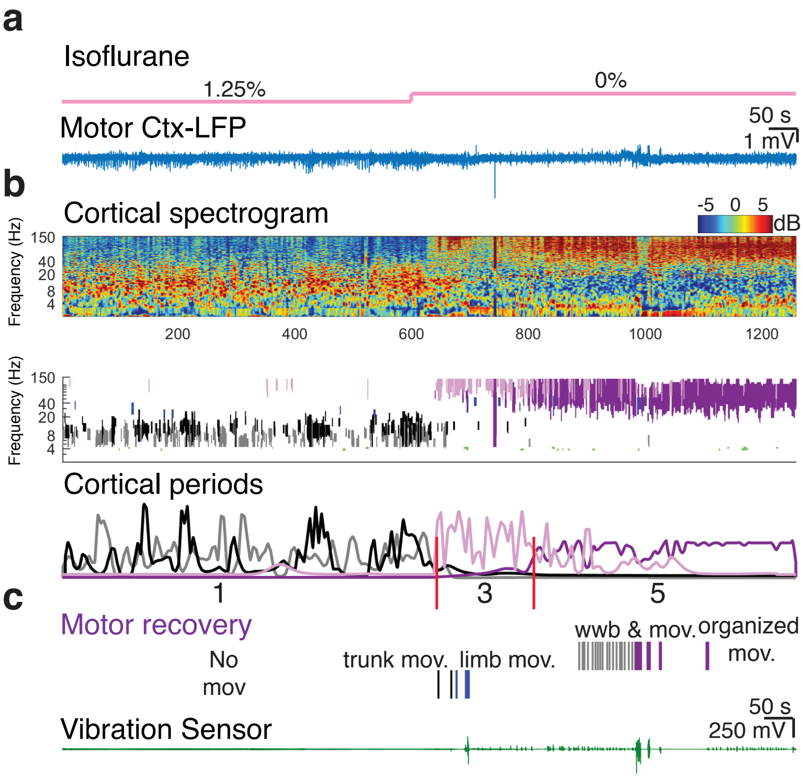


**Supplementary Figure 2**
